# Antiviral activity versus host response: contrasting transcriptomic footprints of four antivirals in human cytomegalovirus-infected brain organoids

**DOI:** 10.64898/2026.05.01.722178

**Authors:** Ece Egilmezer, William Rawlinson, Charles S.P. Foster

**Affiliations:** Kirby Institute, University of New South Wales, Sydney, NSW, Australia; Serology and Virology Division (SAViD), NSW Health Pathology, Prince of Wales Hospital, Sydney, NSW 2031, Australia; School of Biomedical Sciences, Faculty of Medicine & Health, University of New South Wales, Sydney, NSW 2052, Australia; School of Biotechnology and Biomolecular Sciences, Faculty of Science, University of New South Wales, Sydney, NSW 2052, Australia

**Keywords:** human cytomegalovirus, cerebral organoids, antiviral therapy

## Abstract

Infection with human cytomegalovirus (HCMV) is common and usually asymptomatic in healthy individuals, but can cause severe neurological injury, particularly following congenital transmission. For symptomatic congenital infection, standard antiviral treatment is ganciclovir, with maribavir and letermovir as alternative direct-acting agents. However, their relative antiviral activity in reducing HCMV RNA abundance and restoring the host transcriptome towards an uninfected state has not been directly assessed in a neural model. We infected human cerebral organoids with Merlin-strain HCMV and treated them for 14 days with aciclovir, ganciclovir, letermovir, or maribavir, comparing each with untreated infected organoids (NO). Relative to NO, antiviral treatment reduced viral load 3.3-fold with aciclovir, 20.1-fold with ganciclovir, 65.4-fold with letermovir, and 6.9-fold with maribavir. We identified >2,500 differentially expressed host genes in infected versus uninfected organoids, with enrichment of neurodevelopmental and metabolic stress pathways. Aciclovir, ganciclovir, and maribavir produced few differentially expressed host genes relative to NO and no significant GO or KEGG enrichment. In contrast, letermovir altered 312 genes enriched for glycolysis and related metabolic processes. An MSigDB Hallmark pathway analysis showed minimal perturbation with aciclovir and letermovir, whereas ganciclovir and maribavir produced small, coordinated pathway-level shifts. This was partly in the same direction as control uninfected organoids but also with additional perturbations not seen in controls. Collectively, antiviral choice influenced both the magnitude of reduction in HCMV RNA abundance and the gene expression profile of infected neural tissue. The results support further evaluation of ganciclovir and letermovir in therapy of neural damage resulting from HCMV infection.

## Introduction

Human cytomegalovirus (HCMV; *Betaherpesvirinae*: *Cytomegalovirus humanbeta5*) is a double-stranded DNA virus with a large ∼235,000 nt genome encoding >750 open reading frames, of which ∼200 encode functional genes [1]. Seroprevalence rates of HCMV are high worldwide, ranging from 80–100%, with slightly higher rates among women of reproductive age [2,3]. Primary infection of healthy individuals with HCMV is usually minimally symptomatic, with reinfection and reactivation being asymptomatic [3]. Individuals with immature (fetus) or compromised immune systems such as organ transplant recipients may develop serious systemic and neuronal pathology [4,5]. HCMV is the leading non-genetic cause of congenital malformation in developed countries, causing significant morbidity and mortality [6]. In developed countries, the incidence of HCMV births ranges from 0.2% to 2.0% and in developing countries, the incidence ranges from 0.6% to 6.1% [7]. Even among infants asymptomatic at birth, congenital HCMV can lead to severe later disease, including central nervous system (CNS) abnormalities (e.g. microcephaly, intracranial calcifications), seizures, neurodevelopmental impairments such as cerebral palsy, and sensory deficits (e.g. frequently sensorineural hearing loss, less frequently visual impairment) [8–10].

The standard of care for symptomatic HCMV infection and prophylaxis has long been treatment with the nucleoside analogue ganciclovir or its prodrug valganciclovir [11,12]. The critical ethical constraints of therapeutic use of such antivirals in congenital infections include (i) exclusion of pregnant women and human fetuses by TGA and FDA regulations, (ii) the host-specific nature of most congenital infections, rendering animal models inaccurate, (iii) the lack of satisfactory animal models for mother to child transfer (MTCT), and (iv) the lack of organ cross-talk in whole tissue models, all of which ignore the complexity of HCMV-host cell interaction [13]. Traditional therapeutics for congenital infections are not used during pregnancy mainly due to drug toxicity.

For example, ganciclovir is limited by significant toxicities, including neutropenia [14], and resistance can emerge during prolonged treatment or prophylaxis [15–17], requiring monitoring for resistance-associated mutations [17]. Despite its success against other herpesviruses (HSV-1 and HSV-2), the nucleoside analogue aciclovir has only limited efficacy against HCMV [18].

In response to efficacy and tolerability concerns, several direct-acting antivirals have been developed for treatment and prophylaxis of HCMV infection. Maribavir is increasingly used in patients who are unable to tolerate ganciclovir or who harbour ganciclovir-resistant virus [19]. Maribavir acts by directly inhibiting the HCMV serine/threonine protein kinase UL97 [20]. Although early clinical trials yielded mixed results [21], more recent studies have demonstrated the efficacy of maribavir for both prophylaxis and treatment of viraemia [22]. Moreover, it is generally better tolerated than ganciclovir and foscarnet, with substantially lower rates of neutropenia and acute kidney injury, respectively [19]. Letermovir is a more recently developed antiviral that targets the HCMV terminase complex and blocks late genome packaging [23]. It is most commonly used as prophylaxis [24], with ganciclovir or maribavir typically preferred for established viraemia, and its main adverse effects are gastrointestinal, with serious organ toxicity relatively uncommon [25].

HCMV infection profoundly remodels the transcriptomic landscape of the developing brain, driving dysregulation of cellular metabolism and suppression of key neurodevelopmental programmes [26,27]. Consequently, antiviral therapy should not only achieve viral clearance or durable suppression, but also provide an opportunity for endogenous tissue repair processes and neurodevelopmental gene-expression programmes to begin restoring normal brain structure and function. For example, cerebral organoid models of HSV-1 encephalitis have shown that aciclovir can potently inhibit viral replication but leave widespread inflammatory gene-expression responses largely unresolved, highlighting that suppression of viral replication does not necessarily allow host tissue states to normalise [28]. The capacity of current HCMV antivirals, including ganciclovir, maribavir, and letermovir, to shift host gene-expression profiles toward an uninfected reference state in neural tissues remains largely unknown. In principle, suppressing viral replication should eliminate the main driver of dysregulation and allow host cells to recover broad transcriptional homeostasis. However, this recovery is not guaranteed because distinct antiviral mechanisms - differentially targeting different stages of the viral life cycle - may produce distinct host gene-expression signatures and/or induce specific stress responses that prevent a full return to an uninfected baseline.

Organoids derived from human pluripotent stem or progenitor cells are tissue models that mimic human organ development, and have been established to differentiate into a number of different organ types including the brain [29]. Since establishment of cerebral organoid generation protocols, these tissue models have been used to model neurodevelopmental phenotypes arising from viral infection such as HCMV and Zika virus [26,30]. Previously, we demonstrated that HCMV infection of human cerebral organoids dysregulated critical neurodevelopmental pathways (including neuronal pluripotency and differentiation pathways). We showed modifiable pathogenetic mechanisms for HCMV-induced neural malformation and associated development of ASD-like phenotypes [26]. Another recent study using iPSC-derived human neural cultures (2D neural progenitor cells and dorsal forebrain regionalised organoids) showed ganciclovir, letermovir, and maribavir all effectively limited HCMV replication [31]. Some evidence of organoid host gene expression restoration was observed, but only seven host genes were measured. Therefore, it remains critical to determine whether these antivirals are equivalent in their ability to normalise brain-relevant gene expression at the pathway level post-HCMV clearance, or whether their distinct mechanisms of action are associated with different effects on virus-induced host gene expression.

Here, we quantitatively investigated the transcriptome-wide effects of aciclovir, ganciclovir, letermovir, and maribavir in HCMV-infected cerebral organoids, examining both virus and host gene expression. We specifically examined virological suppression alongside treatment-associated host gene-expression responses in a model with complex cytoarchitecture and active early brain developmental programs [29]. We found that, although all antivirals reduced viral load, the magnitude of reduction differed among treatments and they drove distinct and often incomplete trajectories of host gene expression restoration towards an uninfected baseline state. These findings show how the course of gene expression changes in HCMV-infected neural tissues may be differentially shaped by antiviral choice, especially with respect to metabolic programmes.

## Results

### Cerebral organoids generated from human iPSCs display markers for cerebral differentiation

The HCMV infection and antiviral responses were evaluated in a human-relevant developing brain context. We generated 15 cerebral organoids from iPSCs and confirmed their regional identity and cellular composition. Organoids cultured for 55 days showed cerebral differentiation consistent with our previous work [26]. They expressed Nestin (neural progenitor cells) and βIII-tubulin (neurons), indicating neuronal differentiation, and contained GFAP-positive regions consistent with astrocyte formation. Regionally, FOXG1 staining supported forebrain identity, while TBR1 positivity indicated pre-plate neuron generation (data not shown).

### RNAseq overview and data exploration

We generated bulk RNA-seq data from the 15 organoids that were infected with human cytomegalovirus (HCMV). Identical numbers of neuronal organoids were either left untreated (“NO”, *n* = 3) or treated with different antivirals, including aciclovir (“ACV”, *n* = 3), ganciclovir (“GCV”, *n* = 3), letermovir (“LMV”, *n* = 3), and maribavir (“MBV”, *n* = 3). We additionally included comparator bulk sequencing data from our previous study (“Study 1”, [26]) comprising untreated HCMV-infected (“NO”, *n* = 4) and mock-infected brain organoids (“UNINFECTED”, *n* = 4).

Uninfected samples were presented for quality control (QC) and for contextual reference to understand the expected differences in gene expression relative to an infected, untreated brain organoid. Antiviral-treated and uninfected organoids were not directly contrasted because antiviral treatment status was fully confounded with study batch, as outlined in Methods, and instead an overlap-based approach was used to assess “restoration” to baseline host gene expression (see below).

The expression of HCMV genes and host (human) reads was jointly quantified using a combined reference comprising the human and HCMV genomes. Principal components analysis (PCA) of batch-adjusted log-counts per million expression data recovered no clean separation of treatment groups when focusing on all host genes (Figure 1a), with only PC2 (13.3%) separating uninfected samples from all infected samples. Restricting the PCA to host genes identified as differentially expressed across conditions (see differential expression analyses below) revealed clear clustering by treatment group along PC1 (49.1%), with infected vs uninfected samples separated primarily along PC2 (20.3%) (Figure 1b). Treatment-associated structure was even clearer based on HCMV gene expression: each set of samples corresponding to antiviral treatments formed its own cohesive group and PC1 explained nearly all of the variance (97.1%) (Figure 1c). The GCV and LMV treatments were most distinct from the other treatment groups and differed from one another along PC2, albeit with this axis explaining only a small proportion of the variance (1.7%).

**Figure 1:**
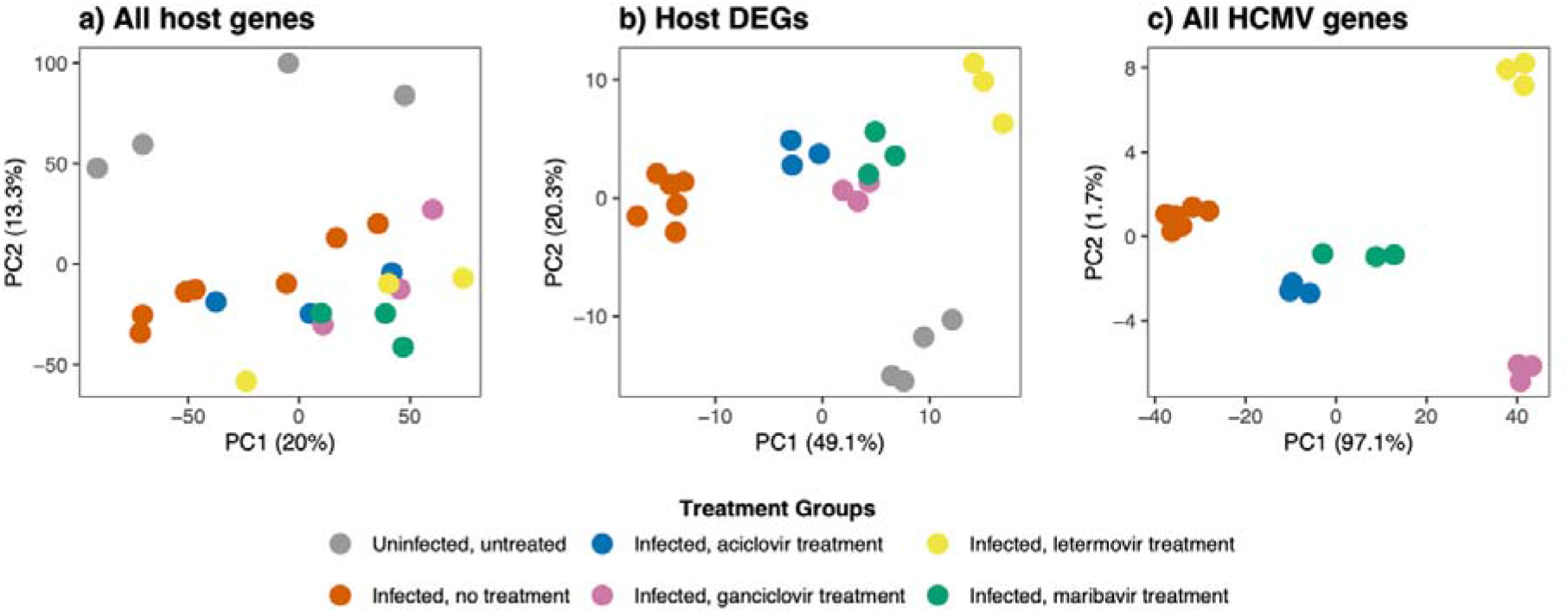
Principal component analysis (PCA) of host and viral transcriptomes across infection and antiviral treatments. PCA plots show clustering of cerebral organoid samples based on (a) all detected human host genes, (b) human host differentially expressed genes (DEGs; defined in Methods), and (c) all detected HCMV genes. Each point represents one organoid replicate; colours indicate treatment group (uninfected/untreated; infected/no treatment; infected + aciclovir, ganciclovir, letermovir, or maribavir). Percent variance explained by each component is shown on the axes. Note: panel (b) is included as a descriptive post-selection PCA rather than independent evidence of group separation.

Given the inherent variability of *in vitro* brain organoid cultures [26,29], we performed reference-based cell-type deconvolution as an orthogonal quality-control check to confirm that each sample reflected the expected mix of canonical cerebral organoid cell states. Using excitatory neurons, inhibitory neurons, intermediate progenitor cells, radial glia, and glycolysis-associated cells as reference categories, we observed broadly similar inferred cellular compositions across treatment groups (Figure 2; Supplementary Table S1). One notable exception was the NO samples from Study 1, for which the estimated inhibitory neuron fraction was ∼0 across samples (Figure 2a). Although several treatment–NO contrasts reached significance after multiple-testing correction, the corresponding effect sizes were small. All estimated shifts were within ±10% and most within ±5% absolute change in proportion (Figure 2b; Supplementary Table S2).

**Figure 2:**
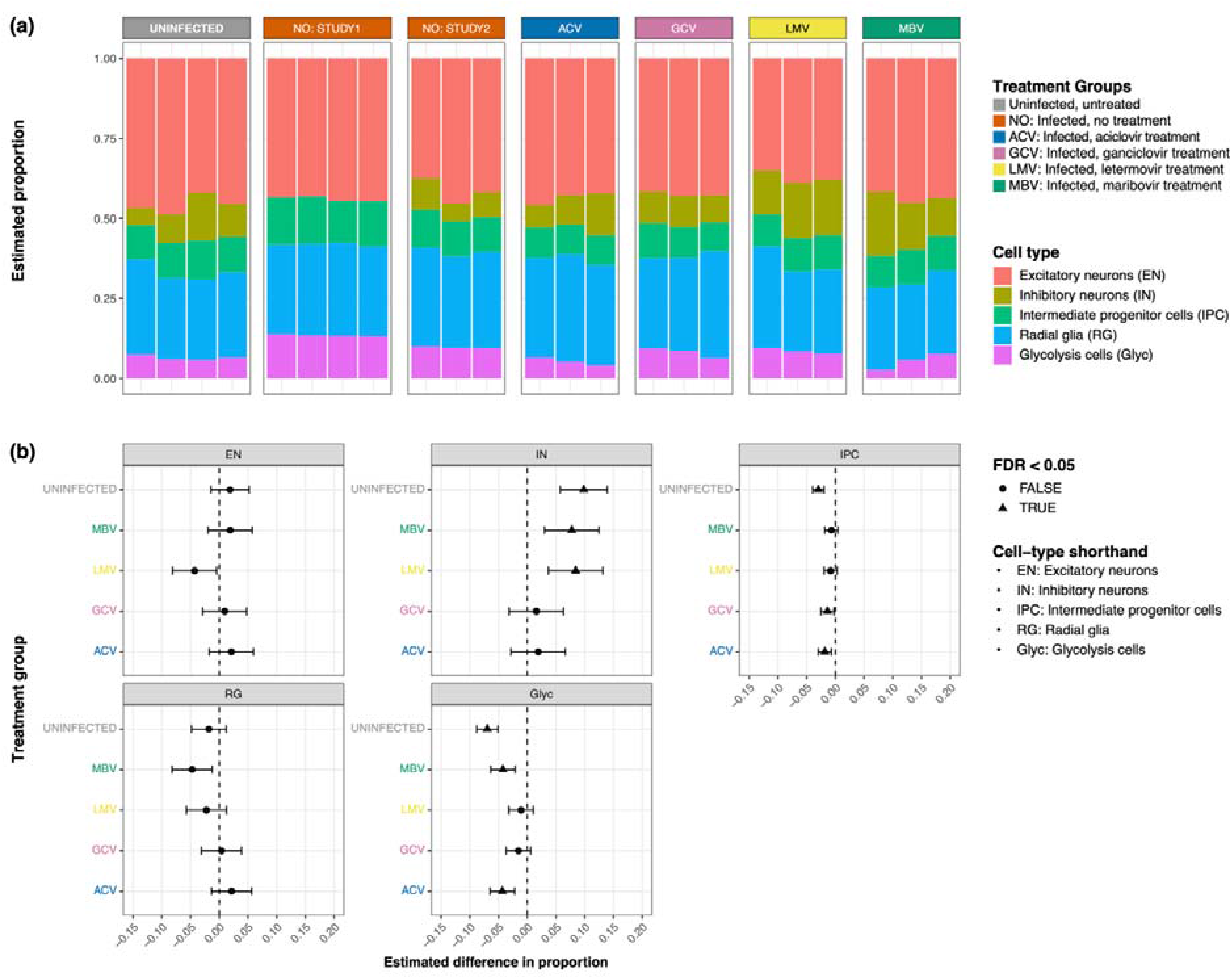
Bulk deconvolution analyses to infer cerebral organoid cell-type compositions. (a) Stacked barplots show the estimated proportions of five broad cell states (excitatory neurons [EN], inhibitory neurons [IN], intermediate progenitor cells [IPC], radial glia [RG], and a glycolysis-associated state [Glyc]) for each sample, grouped by condition (uninfected/untreated; infected/no treatment across two batches/studies; infected + aciclovir [ACV], ganciclovir [GCV], letermovir [LMV], or maribavir [MBV]). (b) Estimated differences in cell-type proportion for each treatment group relative to the infected, no-treatment condition are shown with confidence intervals; triangles denote contrasts significant at FDR < 0.05.

### Fixed-concentration antiviral treatments differ in their reduction of HCMV viral load

To compare antiviral activity at the fixed concentration tested, we estimated the log -fold reduction in normalised HCMV-aligned RNA-seq abundance in each antiviral-treated group relative to the infected, untreated (NO) group, used here as a proxy for tissue-level viral load. This measure reflects aggregate viral RNA abundance and does not directly quantify viral genome copies or infectious virus. All four antivirals resulted in a reduction in the HCMV viral load, but with the magnitude of success varying among groups (Figure 3; Supplementary Table S3). The smallest reduction (3.3-fold) in HCMV viral load was in aciclovir-treated organoids, followed by maribavir (6.9-fold reduction). Ganciclovir produced a substantial reduction in HCMV viral load (20.1-fold; 95% reduction), while letermovir achieved the greatest effect (65.4-fold; 98.5% reduction).

**Figure 3.**
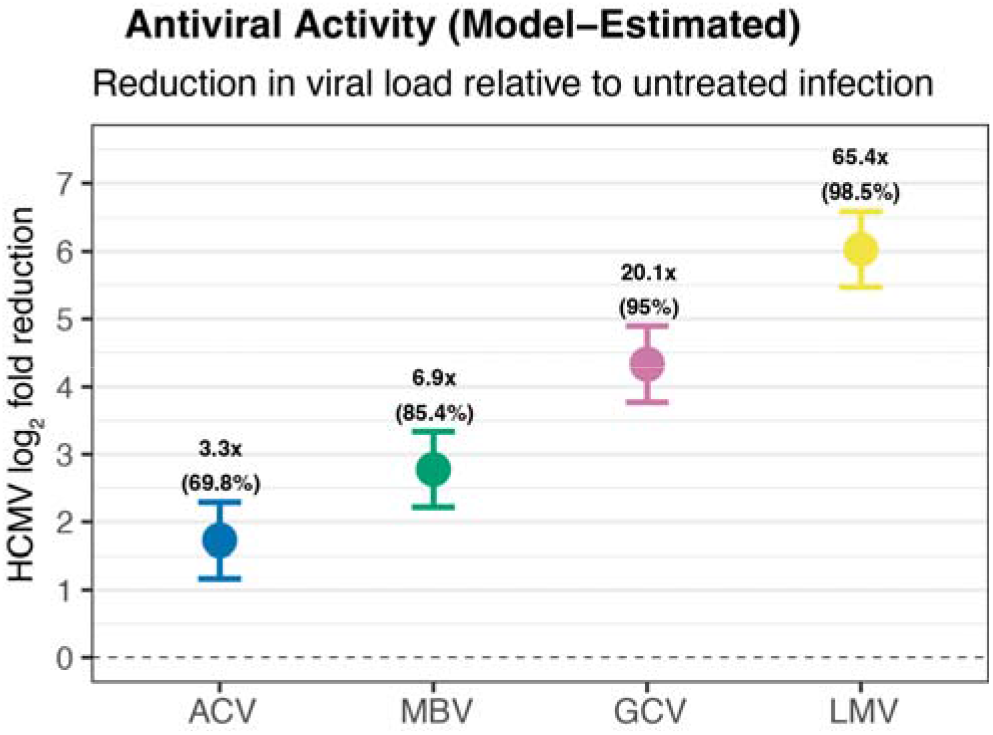
Model-estimated reductions in HCMV viral load in HCMV-infected cerebral organoids following antiviral treatment. Points show the estimated reduction in HCMV viral load (log_2_ fold change) for each antiviral treatment relative to infected, untreated controls; error bars indicate uncertainty (95% CI). Numbers above points report the corresponding fold reduction in viral load and the percent reduction. Dashed line marks no change (0). ACV, aciclovir; MBV, maribavir; GCV, ganciclovir; LMV, letermovir.

Together with similar total and host-aligned read depths across groups, these results indicate that the markedly lower HCMV read fractions in LMV-treated organoids reflect genuine virological suppression rather than differences in sequencing depth or host RNA content. Consistent with this, qPCR quantification of HCMV major immediate-early (MIE) signal showed significant reductions in DNA abundance for all antivirals relative to NO (GCV p<0.0001, LMV p<0.0001, MBV p<0.0001, ACV p=0.0002), providing orthogonal support for the observed antiviral activity (Figure S1).

Although ganciclovir and letermovir ranked differently by RNA-seq and MIE-DNA qPCR, both assays supported substantial antiviral activity, with the difference potentially reflecting their measurement of distinct viral nucleic-acid endpoints.

Out of the 155 HCMV genes passing filtering thresholds, 154 were found to be differentially expressed in contrasts between each antiviral group and NO (Figure 4; Supplementary Table S4). The only gene not differentially expressed in each case was US34. The observed patterns indicate a largely global reduction in viral gene expression, rather than selective targeting of specific kinetic classes, and we did not observe clear differential targeting of any of the seven temporal classes of HCMV genes defined by Rozman et al. [32]. To exclude confounding by pre-existing or emergent antiviral resistance in our HCMV inoculum, we analysed all non-human reads per sample and called HCMV variants against the Merlin genome. No known resistance-associated mutations were detected in any treatment group, indicating that antiviral resistance is unlikely to explain the observed differences in virological signals and host gene-expression profiles. The observed patterns are consistent with contributions from differences in drug mechanism and potency; however, these effects cannot currently be separated from differences in residual viral burden or direct drug effects. Only one non-synonymous mutation was detected (C11356CA), with the inferred impact of a frameshift in the *RL13* gene, as was expected given that we used an *RL13*- mutant stock of HCMV.

**Figure 4.**
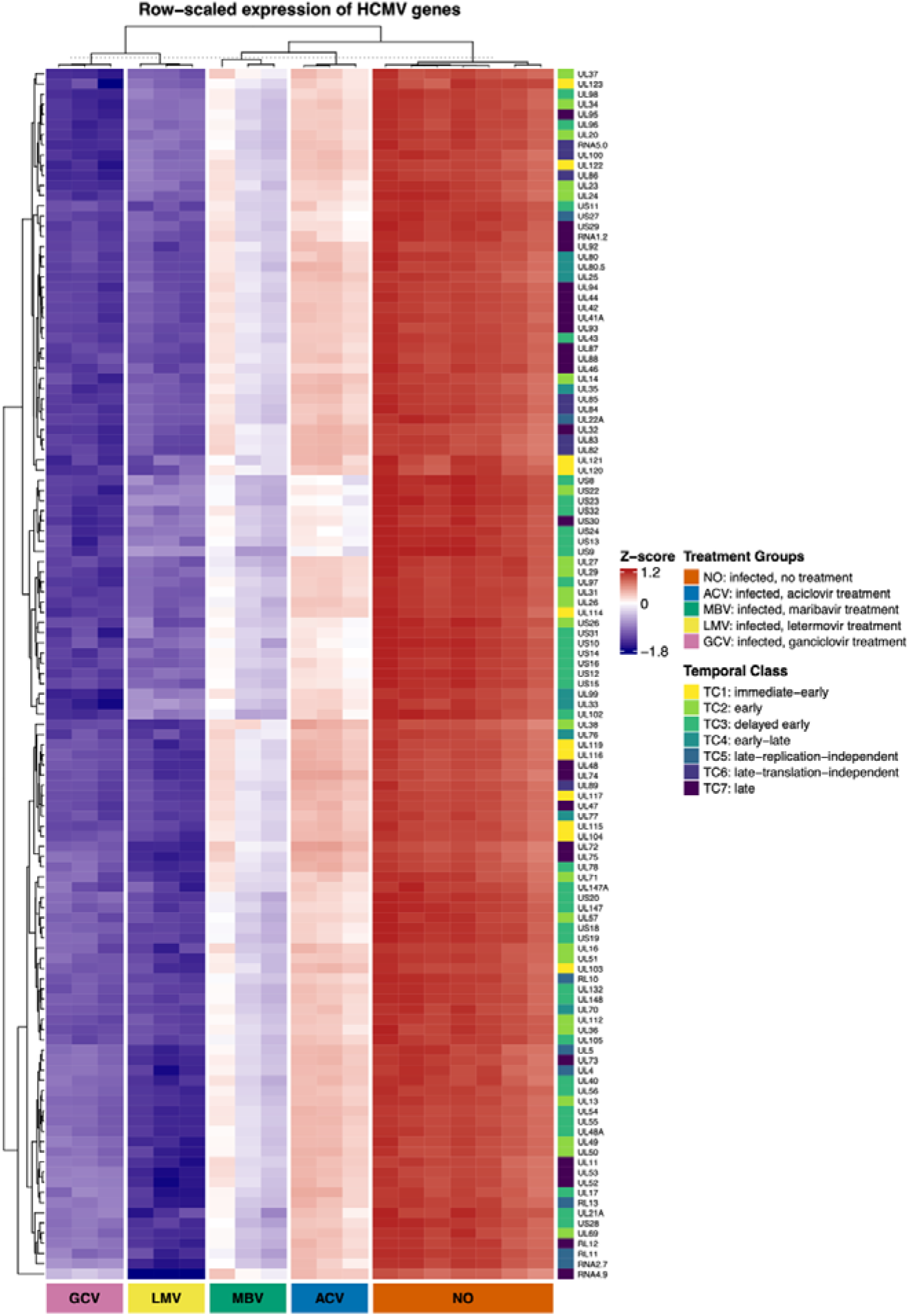
Antiviral treatments produce distinct HCMV RNA expression profiles in infected cerebral organoids. Heatmap shows row-scaled (Z-score) expression of HCMV genes across infected organoid samples by treatment group (NO = infected, no treatment; ACV = aciclovir; GCV = ganciclovir; LMV = letermovir; MBV = maribavir). Columns were hierarchically clustered globally (Euclidean distance; complete linkage) and then split by treatment group; rows were hierarchically clustered (Euclidean; complete). The right-hand annotation indicates each gene’s temporal expression class (TC1–TC7) according to the scheme of Rozman et al. [32]. Warmer colours indicate higher relative expression of a given gene across samples; cooler colours indicate lower relative expression.

### Differential host gene expression after antiviral treatments

Our analyses of HCMV gene expression above characterise antiviral activity separately from host gene expression responses, which we assessed by analysing host differentially expressed genes and pathway-level perturbations in the same samples. We previously demonstrated that infection with HCMV results in dysregulation of key host cellular genes and pathways in cerebral organoids [26]. These patterns were recapitulated in our reanalysis of the prior data combined with additional untreated, HCMV-infected brain organoids in the present study. In contrasts between NO and UNINFECTED groups, 2547 host genes were recovered as differentially expressed (Supplementary Table S5). These differentially expressed genes (DEGs) were significantly enriched for 31 KEGG terms including neurological disease, including “pathways of neurodegeneration - multiple diseases” (HSA05022), “spinocerebellar ataxia” (HSA05017), and 402 GO terms that reflected general dysregulation of brain development and function, including “neuron differentiation” (GO:0030182), “regulation of axonogenesis” (GO:0050770), and “brain development” (GO:0007420) (Supplementary Table S6). Competitive gene set testing of MSigDB Hallmark gene sets revealed coordinated directional changes in gene expression between the UNINFECTED and NO groups. Of the 50 Hallmark gene sets, 21 were enriched for differential expression relative to genes outside the sets (Supplementary Table S7). These enriched gene sets represent perturbation of five broad groupings of biological processes including proliferation and DNA repair, metabolism and stress, immune and inflammation, development and structure, and other signalling processes (Figure 5; Supplementary Table S7).

**Figure 5:**
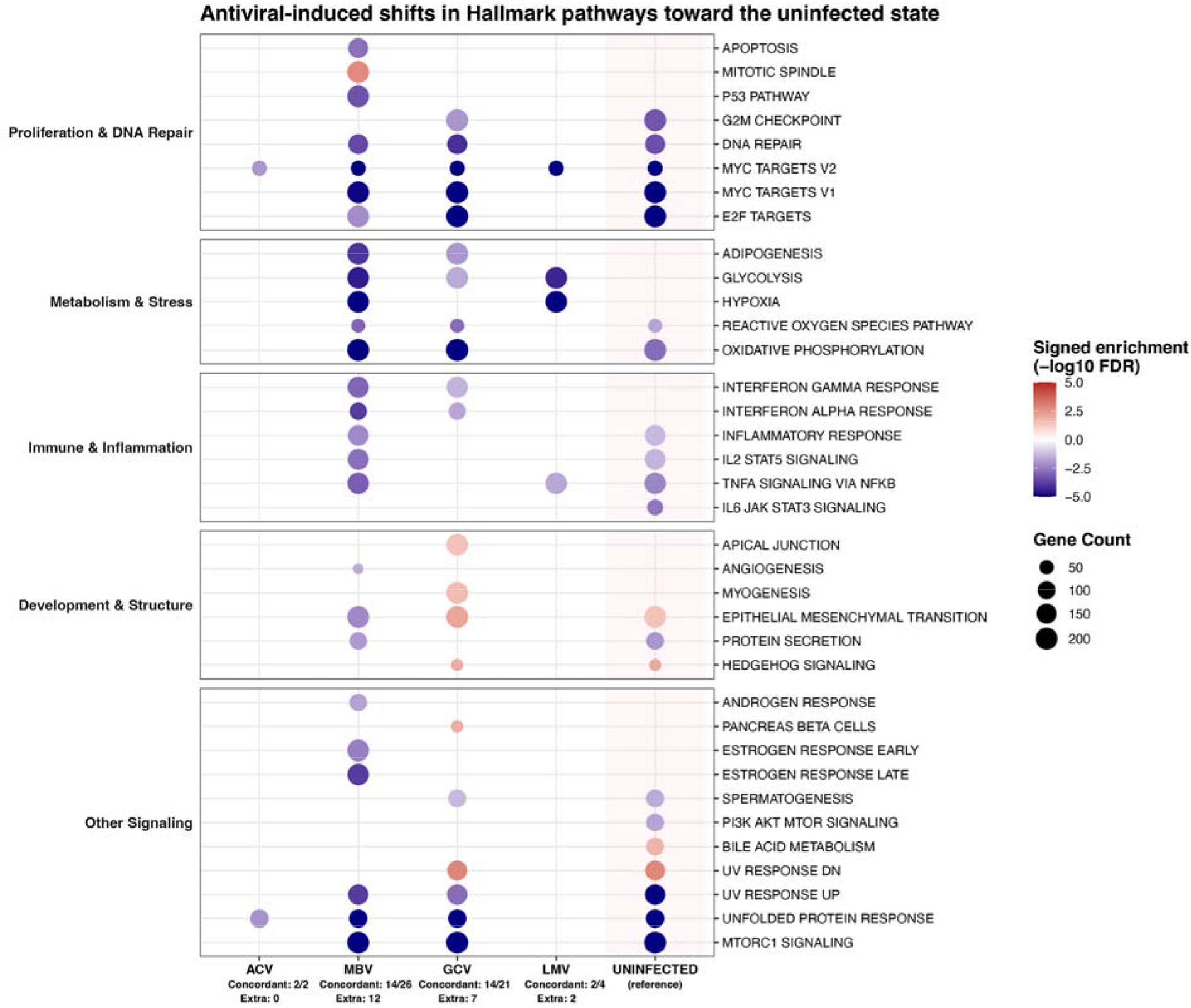
Antiviral-induced perturbation of Hallmark pathways toward the uninfected state. Dot plot summarising Hallmark gene set enrichment changes in infected cerebral organoids treated with aciclovir (ACV), ganciclovir (GCV), letermovir (LMV) or maribavir (MBV), relative to untreated infection, with the uninfected condition shown as the reference. Pathways are grouped into broad functional modules (left). Dot colour indicates signed enrichment (−log10 FDR; red = positive, blue = negative), and dot size indicates the number of genes contributing to each set. Values beneath each treatment indicate the number of pathways directionally concordant with the uninfected-versus-infected reference contrast (“concordant”) and additional significant pathways not classed as concordant (“extra”).

Differential host gene expression between antiviral-treated and the NO groups varied among each antiviral group (Figure 6a; Supplementary Table S5). Relatively few genes were differentially expressed between the NO and ACV, MBV and GCV experimental groups, and these DEGs were not enriched for any GO or KEGG terms (Supplementary Table S6). In contrast, 312 genes were differentially expressed between NO and LMV, and these DEGs were enriched for GO terms and KEGG pathways related to signalling and metabolic processes (Supplementary Tables S5–S6). Most LMV-associated DEGs were unique to LMV, MBV and GCV shared a modest core of host genes with altered expression, and only a small number of genes were perturbed by more than two antivirals (Figure 6b). Because ganciclovir and letermovir produced the largest reductions in HCMV viral load, we directly contrasted the GCV and LMV groups to characterise their gene-expression differences. By doing so, we aimed to address whether letermovir effects resulted from a greater reduction in viral load, or the antiviral induced a host state qualitatively distinct from that of ganciclovir-treated organoids. The 85 DEGs assessed in this direct comparison revealed that, relative to GCV, letermovir-treated organoids showed coordinated downregulation of glycolysis, HIF-1 signalling, and nucleotide metabolic pathways, alongside relative upregulation of aminoacyl– tRNA biosynthesis and unfolded protein response signatures (Supplementary Tables S5–S7).

**Figure 6:**
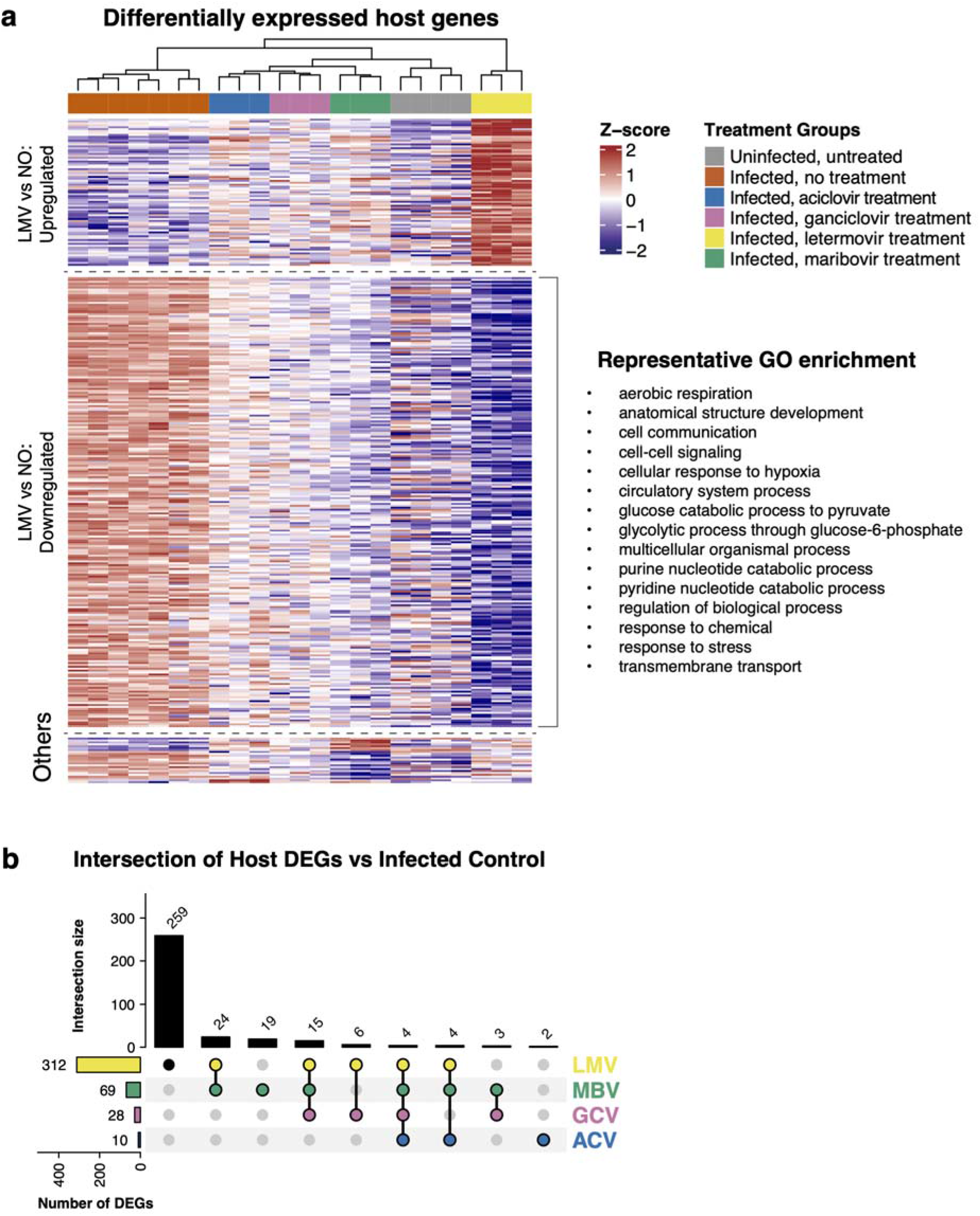
Letermovir treatment produces a distinct host gene expression state relative to untreated infection. (a) Heatmap of host differentially expressed genes (DEGs) across treatment groups (uninfected; infected untreated [“NO”]; and treatment with aciclovir [ACV], ganciclovir [GCV], letermovir [LMV], or maribavir [MBV]). Genes are shown as row-scaled expression (Z-scores) and both rows and columns are hierarchically clustered (Euclidean distance; complete linkage); major gene blocks highlight genes upregulated or downregulated in LMV versus NO, with representative enriched GO terms shown. (b) UpSet plot showing the overlap of host DEGs (each antiviral vs infected untreated control), with bar heights indicating intersection sizes and the left bars showing total DEGs per treatment.

These results are consistent with a shift away from the highly glycolytic, hypoxia-response state observed under ganciclovir treatment of organoids towards a metabolically down-shifted but proteostatically stressed phenotype. Thus, the gene expression changes induced by LMV were distinct in terms of gene function, consistent with the possibility that its mechanism of action produces a host gene expression response differing from that associated with ganciclovir treatment.

Given the demonstrated impact of HCMV infection on brain organoid function, and the clear effectiveness of antiviral treatments in suppressing HCMV in organoids and clinical settings, we investigated whether treatment with antivirals could return host gene expression towards the healthy baseline. We hypothesised that over the time frame of our experiment we would not find a total return to baseline host gene expression, but rather small and coordinated shifts in the expression of genes in pathways that might suggest a “partial rescue” effect. Competitive gene set testing supported this hypothesis and revealed qualitatively different impacts of treatment with each antiviral on host gene expression. Only a minority of the 50 Hallmark gene sets were significantly perturbed in the ACV (two perturbed) and LMV (four perturbed) groups, respectively (Supplementary Table S7; Figure 5). In contrast, more gene sets were significantly perturbed in GCV (21 perturbed) and MBV (26 perturbed). Many of these gene sets overlapped directionally with those gene sets enriched in the UNINFECTED vs NO comparison (GCV:14/21; MBV: 14/26), but with additional gene sets enriched in both GCV and MBV respectively (Supplementary Table S7; Figure 5).

## Discussion

Congenital infection with human cytomegalovirus is a significant cause of serious neuropathogenesis in fetal and infant brains [8,33,34]. Our experimental design using a model neuronal organoid system [35] provided a novel means to assess host gene expression changes associated with antiviral-mediated control of HCMV and the subsequent host response. Comparing mock-infected brain organoids with infected organoids, we uncovered many DEGs enriched for GO terms and KEGG pathways related to neurological and metabolic stress, with similar coordinated gene set perturbations (Supplementary Tables S5–7; Figure 5). Crucially, the antiviral treatments differed both in their effects on HCMV RNA abundance and in the extent to which host gene expression shifted towards the uninfected reference profile. These differences may reflect the drugs’ distinct mechanisms of action, although effects of drug identity cannot be separated fully from differences in residual viral burden.

While all tested antivirals significantly reduced HCMV viral load, there was a clear difference in effects among treatments (Figure 3; Supplementary Table S3). The relatively smaller reduction in HCMV-aligned RNA-seq reads following aciclovir treatment relative to untreated infection, our measure of fixed-concentration antiviral activity, is consistent with its limited efficacy against HCMV *in vitro* and *in vivo* [18]. The current reference standard for treatment of symptomatic HCMV infection is ganciclovir [11], with maribavir currently a second-line drug [36]. A recent clinical trial (“AURORA”) demonstrated the superior efficacy of ganciclovir for treatment of highly immune-compromised haemopoietic stem cell transplant patients compared to maribavir, and with more resistance mutations after maribavir treatment [37]. At the fixed concentration tested, our results were directionally consistent with the AURORA clinical trial, with a far larger reduction in HCMV RNA abundance with ganciclovir than maribavir (Figure 3). Although poor blood–brain barrier penetration may limit maribavir activity *in vivo* [19,38,39], this cannot explain its lower effect in our organoid model, where drug exposure is not constrained by a physiological blood–brain barrier.

Our finding that letermovir produced a larger reduction in viral load than ganciclovir at the concentration tested is somewhat surprising. While a prematurely discontinued clinical trial (“SOLACE”) concluded that treatment of HCMV infection with letermovir was safe and well tolerated [40], the drug is currently only FDA-approved for prophylaxis [24]. There are few clinical trial data evaluating letermovir use for treatment rather than prophylaxis. Our neuronal model results suggest that therapeutic usage of letermovir (rather than purely prophylactic usage) warrants further study, and are consistent with the greater antiviral activity of letermovir relative to ganciclovir reported in HCMV-infected iPSC-derived neural cultures and forebrain organoids [31]. An important caveat is that using a common nominal concentration of 25 µM standardised the extracellular concentration and dosing schedule across the four antivirals but did not match them by potency. Differences in intrinsic potency, protein binding, stability, cellular penetration and, where applicable, intracellular activation mean that 25 µM does not represent equivalent pharmacologically active exposure across compounds. The observed differences therefore establish differential suppression of HCMV RNA abundance under the tested conditions, while comparisons at additional concentrations would be required to characterise their relative dose– response profiles. This limitation is particularly relevant to letermovir, for which 25 µM substantially exceeds reported cell-culture EC50 values [31].

We were also interested in whether antiviral treatment shifted host gene expression programmes towards the uninfected reference profile, using the relative directional changes in expression between untreated, mock-infected organoids (UNINFECTED) and untreated, infected organoids (NO) as the baseline signature of HCMV infection. Previous work suggested two weeks of antiviral treatment is unlikely to be sufficient for full recovery of host programs. For example, in transplant recipients, CMV-specific T cell immunity often reconstitutes only after ∼100 days or more despite effective antiviral control of viraemia [41]. We therefore expected any restorative effects to manifest as partial, pathway-level gene expression shifts rather than complete convergence with the uninfected reference profile, as we observed in the neuronal model.

The suboptimal clearance of HCMV by aciclovir observed *in vivo* and *in vitro* was reflected in the host gene expression results. Relative to NO, aciclovir treatment showed little evidence of transcriptomic recovery within the organoids. There were only 10 genes differentially expressed, with no enriched GO terms or KEGG pathways and only two non-specific Hallmark gene sets (Supplementary Tables S5–7). By contrast, letermovir produced the largest reduction in HCMV RNA abundance and the largest host gene expression shift relative to NO (Figure 3; Figure 6), including coordinated down-regulation of glycolysis, oxidative phosphorylation, and ATP metabolic pathways (Supplementary Tables S3–7). This combination is consistent with the possibility that inhibition of viral genome packaging has consequences for the host expression state that differ from those produced by inhibitors acting at earlier stages of replication. Relative to aciclovir, maribavir and ganciclovir were more effective at reducing HCMV RNA abundance, but this did not translate into full host transcriptome recovery over the experimental time course. Instead, competitive gene set testing showed small but significant shifts in Hallmark gene sets in the same direction as UNINFECTED versus NO, indicating persistent infection-associated inflammation and stress despite partial recovery (Figure 5). Direct comparison of letermovir- and ganciclovir-treated infected organoids further showed that these drugs differed not only in the magnitude of viral-load reduction, but in the host state they left behind (Supplementary Tables S5–7). Relative to ganciclovir, letermovir was associated with a metabolically down-shifted but proteostatically stressed state, consistent with a model in which ganciclovir maintains a glycolytic, hypoxia-like programme whereas letermovir reduces viral metabolic demand but leaves a burden of accumulated viral proteins.

The host gene expression differences associated with each antiviral are consistent with contributions from their distinct mechanisms of action, although differences in residual viral burden and direct drug effects may also contribute. Letermovir inhibits the HCMV terminase complex and blocks late genome packaging [23]. In our data, this late-stage inhibition was associated with broad metabolic down-shifting without restoration of HCMV-perturbed neurodevelopmental pathways.

Specifically, glycolysis, oxidative phosphorylation, and ATP metabolic pathways were coordinately downregulated, driven by reduced expression of key glycolytic enzymes and transporters including HK2, PFKP, PFKFB3, ENO1, PGAM1, GPI, SLC2A1, SLC2A3, and SLC16A3 (Supplementary Table S5). This pattern is consistent with reversal of the Warburg-like metabolic shift that occurs in HCMV replication within cells *in vitro* [42–44]. However, genes involved in neuronal structure and function were also downregulated (e.g. NEFH, SLC6A3, SYNDIG1L, ICAM5; Supplementary Table S5), suggesting that metabolic suppression occurred alongside broader dampening of neurodevelopmental programmes. These findings broadly align with Mulder et al. [31], who reported that letermovir more effectively shifted the expression of selected transcripts (SOX2, PPAR-γ and IL6) towards control levels than ganciclovir in iPSC-derived neural progenitors and dorsal forebrain organoids, but reduced ATP levels and organoid viability at higher doses. Although those specific transcripts were not significantly differentially expressed here, likely due to differences in model system, experimental conditions, and assay, our results likewise indicate that letermovir treatment was associated with a metabolically quiescent, developmentally blunted state that is distinct from both untreated infection and the uninfected baseline, rather than simply shifting host gene expression towards the uninfected reference profile.

In contrast, the gene expression patterns seen under ganciclovir and maribavir are consistent with their earlier and more pleiotropic points of action in the HCMV replication cycle [11,36]. While both drugs ultimately limit productive replication by interfering with viral DNA synthesis and late gene expression, they do so while viral genomes and proteins are still present and sensed by the host. This distinction is relevant because ganciclovir requires UL97-dependent activation before inhibiting viral DNA synthesis, whereas maribavir directly inhibits UL97 kinase activity, a broader regulatory node in HCMV replication [45,46]. The broadest pattern of Hallmark perturbation observed in ganciclovir- and maribavir-treated organoids - encompassing coordinated changes in interferon and TNF/NF-κB signalling, IL2/STAT5 and IL6/JAK/STAT3 pathways, and PI3K–AKT–mTOR and oxidative stress programmes - is consistent with an actively remodelled antiviral and inflammatory state coupled with metabolic down-shifting within the cell. At the gene level, both drugs were associated with down-regulation of neuronal markers (*NEFH*, *SLC6A3*, *GABRQ*, *GIPC3*) and developmental transcription factors (*HES2*, *FOXE1*, *NKX3-1*, *ISL2*, *BARX1*). This is consistent with a partial pathway-level shift towards the uninfected Hallmark profile following antiviral treatment, with neurodevelopmental transcriptional programs remaining perturbed.

Maribavir produced the largest number of “extra” Hallmark changes not seen in uninfected versus untreated comparisons, consistent with the wider role of the virus protein kinase p*UL97* in coordinating late gene expression and virion egress. The changes included additional inflammatory and stress-response signatures, suggesting that UL97 inhibition generated a distinct host response state in which virus replication was curtailed but host signalling networks were still substantially remodelled.

Effective HCMV therapy aims to reduce virus replication and improve long-term clinical neurodevelopmental outcomes [47]. Our results suggest that an ideal HCMV therapeutic would combine robust viral suppression with minimal persistence or induction of abnormal host gene expression programmes. In the case of a developing brain, represented in our brain organoid system, antivirals that both suppress virus and minimise off-target effects are preferable. The clear gene expression differences between letermovir- and ganciclovir-treated organoids, despite substantial reductions in HCMV burden with both drugs, support the hypothesis that targeting late genome packaging rather than viral DNA synthesis may influence not only the amount of virus present but also the gene expression and metabolic state that persists during antiviral treatment. This suggests that antiviral choice could differentially shape the trajectory of neural recovery even when virological endpoints (viral load) appear comparable. However, the absence of uninfected drug-treated organoids and a matched vehicle control prevents separation of infection-dependent responses from direct drug or vehicle effects, and these data do not establish whether treatment-associated gene-expression changes reflect cytotoxicity. The cerebral organoid model used here did not include microglia or infiltrating immune cells and therefore could not capture their contributions to treatment responses. Furthermore, since bulk RNA sequencing averages signals across heterogeneous cell populations, it cannot distinguish reduced viral RNA within infected cells from changes in the number, identity, or distribution of infected cells. Future studies using single-cell or spatial transcriptomics could resolve cell-type-specific host responses and antiviral effects while enabling direct mapping of viral tropism [48].

## Materials and Methods

### Cell lines and preparation of virus stocks

Human episomal induced pluripotent stem cells (iPSC) were cultured as previously described [26]. Briefly, human episomal iPSCs (Gibco) with a normal karyotype were cultured on 6-well plates coated with vitronectin (Gibco) as described by the manufacturers’ protocol. The media used was complete Essential 8 Medium (Gibco) and supplemented with 1X Revitacell (Gibco) for the first 24 hours of passage. For continued iPSC passage and maintenance, the cells were cultured on 6-well plates coated with Growth Factor Reduced Basement Membrane Matrix Matrigel (Corning) [49].

The cells were cultured in complete mTeSR1 Medium (Stem Cell Technologies) and supplemented with ROCK Inhibitor Y-27632 (Sigma-Aldrich) for the first 24 hours of culture. All cell lines were *Mycoplasma* free and maintained at 37°C with 5% CO_2_. The genetically intact HCMV strain Merlin (UL128+, RL13−) was propagated as previously described and standard plaque assays were used to determine viral titers [50].

### Generation of cerebral organoids

Cerebral organoids were generated from iPSCs as previously described [26,49]. Briefly, 9,000 live iPSCs were seeded in a low-attachment 960-well U-bottom plate and cultured in low bFGF hESC medium supplemented with ROCK inhibitor to generate embryoid bodies for 5–7 days. Rock inhibitor and bFGF were only included for the first four days of culture. Every other day, half the medium was removed and replaced with fresh media. Following embryoid body generation, the clusters were transferred to a low-attachment 24-well plate and cultured for 4–5 days in neural induction medium to generate primitive neuroepithelial cell clusters. Without removing the media present in the well, fresh neural induction medium was added to each well every 48 hours of culture. After culture, the neuroepithelial tissues were embedded in Matrigel droplets and cultured in cerebral organoid differentiation medium (without vitamin A) in Petri dishes. The medium was replaced with fresh cerebral organoid differentiation medium (without vitamin A) following 48 hours of culture. Four days after culture, the organoids were transferred to fresh Petri dishes and cultured in cerebral organoid differentiation medium (containing vitamin A). This was cultured on an orbital shaker, replacing the medium every 3-4 days with fresh medium.

Following 55 days of culture, the organoids were confirmed to have undergone cerebral differentiation via immunofluorescence to detect Nestin, βIII-tubulin, GFAP, FOXG1, and TBR1, as per our previous work [26]. At 55 days of organoid generation, the cerebral organoids were inoculated with 1 × 10^7^ PFU of CMV strain Merlin (yielding 0.1 MOI) and incubated at 37°C with 5% CO_2_ on an orbital shaker. After 24 hours of inoculation, the organoids were transferred to fresh Petri dishes and fresh cerebral organoid media (containing vitamin A) was added. Analytical grade ganciclovir, aciclovir, maribavir and letermovir were obtained from Sigma-Aldrich (St. Louis, Missouri, United States) and MedChemExpress (Monmouth Junction, New Jersey, USA). Stock aliquots of these antiviral compounds were prepared in dimethyl sulfoxide (DMSO) and stored at - 80°C. Each experimental condition comprised three individual organoids derived from a single organoid differentiation batch (n = 3); the experiment was not replicated across independent differentiations. At 1 day post infection (dpi), 25 µM of aciclovir, ganciclovir, maribavir or letermovir were added to the fresh organoid medium, with three organoids per treatment condition. All four compounds were tested at a common nominal concentration of 25 µM as a pragmatic fixed-concentration screen. This provided a standardised nominal extracellular exposure and dosing schedule across treatments while allowing activity to be detected for all four compounds, including the comparatively weak HCMV inhibitor aciclovir. The design was intended to compare treatment-associated virological and host gene-expression effects under a common experimental condition, rather than to match compounds by potency or model clinically equivalent exposures. Infected organoids receiving no antiviral treatment served as controls. Fresh antivirals were included with every organoid media change, every 3–4 days of culture. After 14 days of culture following initial inoculation, the organoids were extracted from Matrigel and harvested.

### Organoid lysing

Following 14 days of infection and antiviral treatment, cerebral organoids were harvested for bulk RNA-sequencing, as previously described [26]. Three organoids per treatment condition (untreated, aciclovir, ganciclovir, maribavir, and letermovir) were harvested individually for bulk RNA sequencing. First, to remove the organoids from the Matrigel matrix, they were each transferred to a 48-well plate and pre-chilled Corning Cell Recovery solution (In Vitro Technologies, Noble Park North, Victoria, Australia) was added. Organoids were gently resuspended by pipetting with a cut 1 mL pipette tip and incubated for 20 min at 4 °C. They were then centrifuged at 200 × g for 30 s at 4 °C and washed twice with ice-cold PBS.

### DNA extraction and qPCR

Total nucleic acid was extracted from lysed cerebral organoids using the MagNA Pure LC Total Nucleic Acid Isolation Kit according to the manufacturer’s protocol (Roche). HCMV major immediate-early (MIE) DNA was quantified by real-time qPCR using a LightCycler 480 instrument (Roche) with KAPA SYBR FAST qPCR mix, following the assay described by Hamilton et al. [51] and originally reported by Hutterer et al. [52]. Reactions were carried out under the following conditions: denaturation at 95 °C for 5 min, followed by 45 cycles of denaturation at 94 °C for 15 s, annealing at 60 °C for 20 s, and elongation at 72 °C for 15 s. The HCMV MIE DNA abundance for each antiviral treatment was compared to untreated infected organoids using a one-way ANOVA followed by Bonferroni’s multiple-comparisons test to correct for multiple testing.

### Bulk RNA sequencing

Once the Matrigel was removed and organoid lysed, RNA extraction was performed using the RNeasy mini kit (Qiagen) as per the manufacturer’s protocol. The purity and rough concentration of RNA extracts was quantified using a NanoDrop spectrophotometer (ThermoFisher). Extracts were then submitted to the Ramaciotti Centre for Genomics for total RNA-sequencing using the Illumina Stranded Total RNA with RiboZero Plus kit and a NextSeq 500 HO 2 × 75 bp flowcell.

### Data pre-processing and quantification

In addition to the novel RNAseq data generated in the present study, sequencing data of four HCMV-infected and four mock-infected brain organoids from our previous study (henceforth: “Study 1”, [26]) were downloaded from the European Nucleotide Archive (accessions: SRR22266557–SRR22266564). All RNAseq data were pre-processed using the nf-core/rnaseq workflow (v3.21.0) [53], utilising reproducible software environments from the Biocontainers project [54]. Briefly, read quality was assessed using FastQC v0.12.1 [55], followed by quality trimming and residual adapter removal with fastp v0.24.0 [56]. Reads were then quasi-mapped using salmon v1.10.3 [57] with the extra arguments "--writeUnmappedNames --seqBias --gcBias -- posBias" against a custom reference genome comprising the GRCh38 human reference and the HCMV Merlin reference (NC_006273.2). This combined reference allowed abundance estimation of both human and HCMV transcripts in each sample. Per-sample sequencing, read-retention, alignment and assignment metrics, together with sample accession numbers, are provided in Supplementary Table S8.

### De-convolution analyses

To evaluate whether differences in cellular composition could confound comparisons between treatment groups, we estimated cell-type proportions from the bulk RNA-seq profiles using reference-based deconvolution. A published single-cell RNA-seq atlas of human cerebral organoids (ArrayExpress accession E-MTAB-7552; [58]) was downloaded and imported into R, and the authors’ cell-type annotations were used to define reference populations representing major organoid cell classes (excitatory neurons, inhibitory neurons, intermediate progenitors, radial glia, and glycolytic/stress-like cells). Cell-type proportions in our bulk RNA-seq samples were then inferred using the single-cell reference and BisqueRNA [59]. Differences in inferred proportions across experimental conditions were assessed using linear models including study as a batch covariate.

### Differential expression and functional enrichment analyses

Transcript-level RNAseq abundance estimates for each sample were imported into R v4.5.2 as a SummarizedExperiment object with associated GenomicRanges metadata using tximeta v1.28.0 [60], then summarised to the gene level. The experimental groups for analysis comprised: (i) infected, untreated organoids (“NO”); (ii) infected organoids treated with aciclovir (“ACV”); (iii) infected organoids treated with ganciclovir (“GCV”); (iv) infected organoids treated with letermovir (“LMV”); and (v) infected organoids treated with maribavir (“MBV”). In addition, mock-infected, untreated organoids (“uninfected”) were included as a reference condition representing baseline host gene expression and inferred cellular states.

Antiviral activity was quantified using linear regression of log -transformed, normalised HCMV-aligned RNA-seq abundance (log2_cmv), with treatment group and study included as fixed effects. HCMV-aligned RNA-seq abundance was used as a proxy for tissue-level viral load (“viral load” hereafter); this measure reflects aggregate viral RNA abundance and does not directly quantify viral genome copies or infectious virus. Infected, untreated organoids (NO) and Study 1 were set as the reference levels, so treatment-group coefficients represented the mean difference in log viral load between each antiviral-treated group and NO, adjusted for study. For reporting, we converted these coefficients to fold-reductions in viral load (2^−β), and to percent reductions using (1 − 1/fold-reduction) × 100. P values for the four comparisons were controlled for multiple testing using the Benjamini–Hochberg false discovery rate. In this study, the magnitude of viral-load reduction was interpreted as antiviral activity at the single nominal concentration tested and was not used to estimate dose–response parameters such as EC50 or EC90.

Downstream differential expression analyses were defined to contrast each antiviral treatment condition (ACV, GCV, LMV, MBV) against infected, untreated organoids (NO), with models including study/batch as a covariate. Host (human) and HCMV genes were analysed separately. Because the uninfected controls were generated and sequenced in the prior study, while all antiviral-treated samples were generated in the present study, treatment status and study batch were completely confounded in antiviral-treated-versus-uninfected comparisons.

Consequently, antiviral-treated groups were not directly contrasted with uninfected controls. Uninfected samples were instead used to characterise the infection-associated reference profile through comparisons with untreated infected organoids and for contextual and quality-control analyses.

Global expression patterns were visualised using principal components analysis of transformed counts (log_2_ counts per million; henceforth: log_2_-CPM). Differential expression was then tested using the glmTreat function of edgeR v4.8.0 [61], applying a minimum effect-size threshold via a thresholded test of the null hypothesis |log_2_ FC| ≤ log_2_(1.1) (i.e., ≤1.1-fold).

Functional enrichment of Gene Ontology terms and Kyoto Encyclopedia of Genes and Genomes (KEGG) pathways among host differentially expressed genes (DEGs) was assessed using the goana and kegga functions of limma v3.66.0 [62], respectively. Competitive gene set tests accounting for inter-gene correlation were performed using camera [63], testing for enrichment of Hallmark gene sets of the Molecular Signatures Database [64]. In all analyses, the threshold for statistical significance was a Benjamini-Hochberg corrected p-value (FDR) <0.05. Visualisation of results was performed using the ComplexHeatmap v2.26 and ggplot2 v4.0.1 packages [65,66].

### HCMV resistance mutation analyses

All HCMV reads from the RNAseq data for each sample were analysed to ensure that our differential expression results were not biased by the presence of putative antiviral resistance mutations in our laboratory stock of Merlin-strain HCMV. Briefly, all non-human reads were first extracted using nohuman v0.5.0 with the HPRC.r2 database [67], with the presence of (only) HCMV confirmed using sylph with the “imgvr_c200_v0.3.0” database [68]. The HCMV reads were then processed using vartracker v2.0.0 [69,70], resulting in predicted amino acid consequences for inferred mutations relative to the HCMV Merlin reference genome.

## Acknowledgments

We thank the University of New South Wales, Sydney, and NSW Health Pathology for their ongoing support, especially Dr Stuart Hamilton for his guidance and expertise. This study was supported in part by grants from the UNSW 3 Rs Grant Scheme (RG192608-A) and the Australia-Germany Joint Research Cooperation Scheme (RG192195).

## Data Availability

Novel sequencing data generated for the purposes of this study have been deposited in the European Nucleotide Archive under the project accession PRJEB106831, with sample accessions outlined in Supplementary Table S8. All code underlying the results and conclusions of this study is available at https://github.com/charlesfoster/hcmv_antiviral_manuscript.

## Supplementary Figures

**Figure S1:**
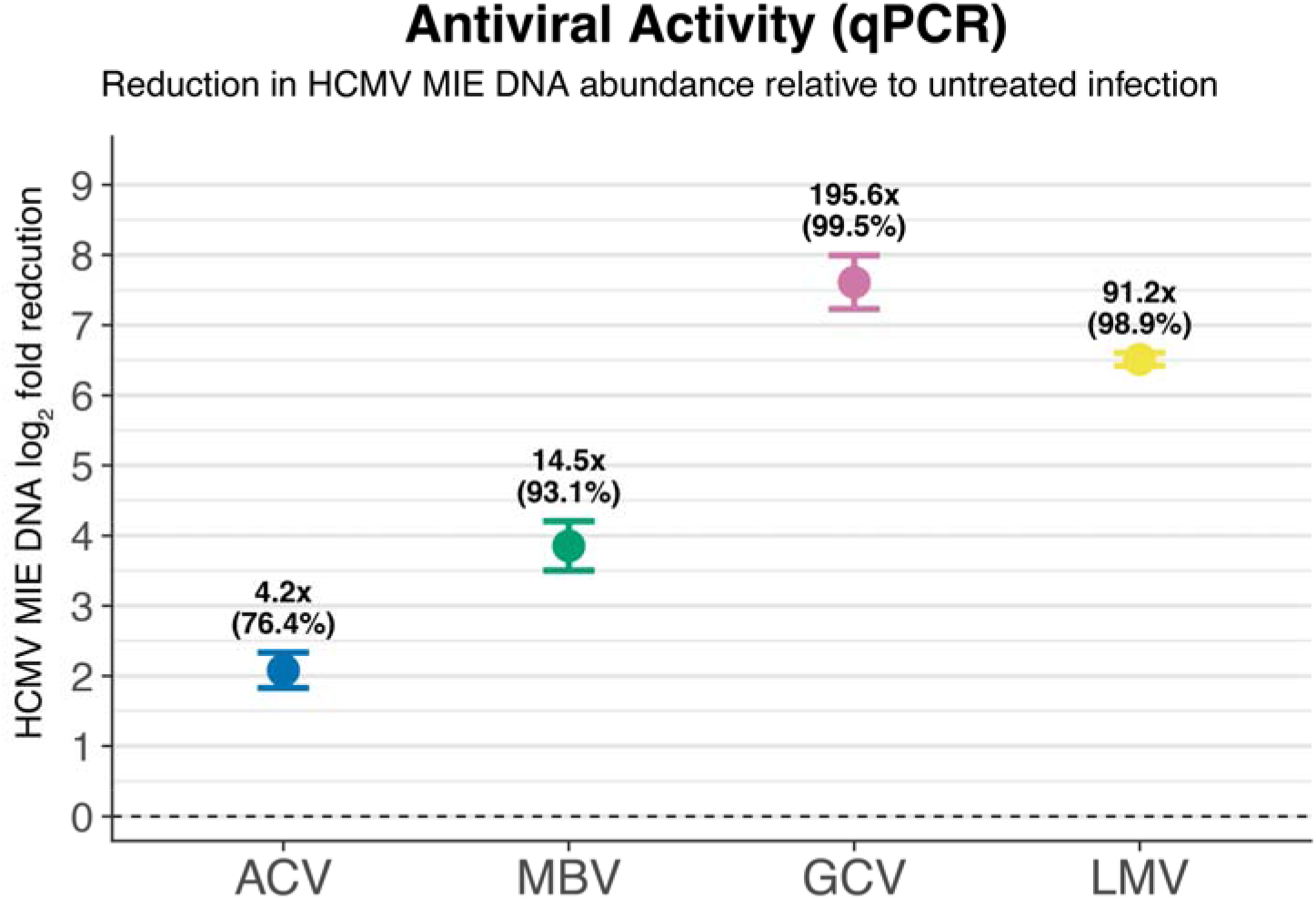
Relative HCMV major immediate-early (MIE) DNA abundance in antiviral-treated, Merlin-infected cerebral organoids. Fifty-five-day-old cerebral organoids were infected with HCMV strain Merlin and treated with antivirals for 14 days. Total nucleic acid was extracted from organoid lysates, and HCMV MIE DNA was measured by real-time qPCR. Values were expressed relative to the untreated infected group, which was set to 100%. Each condition comprised three individual organoids from a single differentiation experiment. Error bars indicate uncertainty (95% CI).

## References

1. Dolan A, Cunningham C, Hector RD, Hassan-Walker AF, Lee L, Addison C, et al. Genetic content of wild-type human cytomegalovirus. Journal of General Virology. 2004;85: 1301– 1312. doi:10.1099/vir.0.79888-0

2. Cannon MJ, Schmid DS, Hyde TB. Review of cytomegalovirus seroprevalence and demographic characteristics associated with infection. Reviews in Medical Virology. 2010;20: 202–213. doi:10.1002/rmv.655

3. Zuhair M, Smit GSA, Wallis G, Jabbar F, Smith C, Devleesschauwer B, et al. Estimation of the worldwide seroprevalence of cytomegalovirus: A systematic review and meta-analysis. Reviews in Medical Virology. 2019;29: e2034. doi:10.1002/rmv.2034

4. Ramanan P, Razonable RR. Cytomegalovirus Infections in Solid Organ Transplantation: A Review. Infection & Chemotherapy. 2013;45: 260–271. doi:10.3947/ic.2013.45.3.260

5. Revello MG, Gerna G. Diagnosis and Management of Human Cytomegalovirus Infection in the Mother, Fetus, and Newborn Infant. Clinical Microbiology Reviews. 2002;15: 680–715. doi:10.1128/cmr.15.4.680-715.2002

6. Grosse SD, Fleming P, Pesch MH, Rawlinson WD. Estimates of congenital cytomegalovirus- attributable infant mortality in high-income countries: A review. Rev Med Virol. 2024;34: e2502. doi:10.1002/rmv.2502

7. Boppana SB, van Boven M, Britt WJ, Gantt S, Griffiths PD, Grosse SD, et al. Vaccine value profile for cytomegalovirus. Vaccine. 2023;41: S53–S75. doi:10.1016/j.vaccine.2023.06.020

8. Cheeran MC-J, Lokensgard JR, Schleiss MR. Neuropathogenesis of Congenital Cytomegalovirus Infection: Disease Mechanisms and Prospects for Intervention. Clinical Microbiology Reviews. 2009;22: 99–126. doi:10.1128/cmr.00023-08

9. Boppana SB, Fowler KB, Vaid Y, Hedlund G, Stagno S, Britt WJ, et al. Neuroradiographic Findings in the Newborn Period and Long-term Outcome in Children With Symptomatic Congenital Cytomegalovirus Infection. Pediatrics. 1997;99: 409–414. doi:10.1542/peds.99.3.409

10. Noyola DE, Demmler GJ, Nelson CT, Griesser C, Williamson WD, Atkins JT, et al. Early predictors of neurodevelopmental outcome in symptomatic congenital cytomegalovirus infection. The Journal of Pediatrics. 2001;138: 325–331. doi:10.1067/mpd.2001.112061

11. Faulds D, Heel RC. Ganciclovir. Drugs. 1990;39: 597–638. doi:10.2165/00003495-199039040-00008

12. Griffiths P, Baraniak I, Reeves M. The pathogenesis of human cytomegalovirus. The Journal of Pathology. 2015;235: 288–297. doi:10.1002/path.4437

13. Schütz M, Cordsmeier A, Wangen C, Horn AHC, Wyler E, Ensser A, et al. The Interactive Complex between Cytomegalovirus Kinase vCDK/pUL97 and Host Factors CDK7–Cyclin H Determines Individual Patterns of Transcription in Infected Cells. Int J Mol Sci. 2023;24: 17421. doi:10.3390/ijms242417421

14. Whitley RJ, Cloud G, Gruber W, Storch GA, Demmler GJ, Jacobs RF, et al. Ganciclovir Treatment of Symptomatic Congenital Cytomegalovirus Infection: Results of a Phase II Study. J Infect Dis. 1997;175: 1080–1086. doi:10.1086/516445

15. Limaye AP, Corey L, Koelle DM, Davis CL, Boeckh M. Emergence of ganciclovir-resistant cytomegalovirus disease among recipients of solid-organ transplants. The Lancet. 2000;356: 645–649. doi:10.1016/S0140-6736(00)02607-6

16. Eid AJ, Arthurs SK, Deziel PJ, Wilhelm MP, Razonable RR. Emergence of drug-resistant cytomegalovirus in the era of valganciclovir prophylaxis: therapeutic implications and outcomes. Clinical Transplantation. 2008;22: 162–170. doi:10.1111/j.1399-0012.2007.00761.x

17. Wong DD, van Zuylen WJ, Novos T, Stocker S, Reuter SE, Au J, et al. Detection of Ganciclovir-Resistant Cytomegalovirus in a Prospective Cohort of Kidney Transplant Recipients Receiving Subtherapeutic Valganciclovir Prophylaxis. Microbiology Spectrum. 2022;10: e02684–21. doi:10.1128/spectrum.02684-21

18. Wagstaff AJ, Faulds D, Goa KL. Aciclovir. Drugs. 1994;47: 153–205. doi:10.2165/00003495-199447010-00009

19. Avery RK, Alain S, Alexander BD, Blumberg EA, Chemaly RF, Cordonnier C, et al. Maribavir for Refractory Cytomegalovirus Infections With or Without Resistance Post-Transplant: Results From a Phase 3 Randomized Clinical Trial. Clin Infect Dis. 2022;75: 690–701. doi:10.1093/cid/ciab988

20. Biron KK, Harvey RJ, Chamberlain SC, Good SS, Smith AA, Davis MG, et al. Potent and Selective Inhibition of Human Cytomegalovirus Replication by 1263W94, a Benzimidazole l- Riboside with a Unique Mode of Action. Antimicrobial Agents and Chemotherapy. 2002;46: 2365–2372. doi:10.1128/aac.46.8.2365-2372.2002

21. Marty FM, Ljungman P, Papanicolaou GA, Winston DJ, Chemaly RF, Strasfeld L, et al. Maribavir prophylaxis for prevention of cytomegalovirus disease in recipients of allogeneic stem-cell transplants: a phase 3, double-blind, placebo-controlled, randomised trial. The Lancet Infectious Diseases. 2011;11: 284–292. doi:10.1016/S1473-3099(11)70024-X

22. Maertens J, Cordonnier C, Jaksch P, Poiré X, Uknis M, Wu J, et al. Maribavir for Preemptive Treatment of Cytomegalovirus Reactivation. New England Journal of Medicine. 2019;381: 1136–1147. doi:10.1056/NEJMoa1714656

23. Goldner T, Hewlett G, Ettischer N, Ruebsamen-Schaeff H, Zimmermann H, Lischka P. The Novel Anticytomegalovirus Compound AIC246 (Letermovir) Inhibits Human Cytomegalovirus Replication through a Specific Antiviral Mechanism That Involves the Viral Terminase. Journal of Virology. 2011;85: 10884–10893. doi:10.1128/jvi.05265-11

24. Imlay HN, Kaul DR. Letermovir and Maribavir for the Treatment and Prevention of Cytomegalovirus Infection in Solid Organ and Stem Cell Transplant Recipients. Clin Infect Dis. 2021;73: 156–160. doi:10.1093/cid/ciaa1713

25. Chemaly RF, Ullmann AJ, Stoelben S, Richard MP, Bornhäuser M, Groth C, et al. Letermovir for Cytomegalovirus Prophylaxis in Hematopoietic-Cell Transplantation. New England Journal of Medicine. 2014;370: 1781–1789. doi:10.1056/NEJMoa1309533

26. Egilmezer E, Hamilton ST, Foster CSP, Marschall M, Rawlinson WD. Human cytomegalovirus (CMV) dysregulates neurodevelopmental pathways in cerebral organoids. Commun Biol. 2024;7: 1–12. doi:10.1038/s42003-024-05923-1

27. O’Brien BS, Mokry RL, Schumacher ML, Pulakanti K, Rao S, Terhune SS, et al. Downregulation of neurodevelopmental gene expression in iPSC-derived cerebral organoids upon infection by human cytomegalovirus. iScience. 2022;25. doi:10.1016/j.isci.2022.104098

28. Rybak-Wolf A, Wyler E, Pentimalli TM, Legnini I, Oliveras Martinez A, Glažar P, et al. Modelling viral encephalitis caused by herpes simplex virus 1 infection in cerebral organoids. Nat Microbiol. 2023;8: 1252–1266. doi:10.1038/s41564-023-01405-y

29. Lancaster MA, Renner M, Martin C-A, Wenzel D, Bicknell LS, Hurles ME, et al. Cerebral organoids model human brain development and microcephaly. Nature. 2013;501: 373–379. doi:10.1038/nature12517

30. Watanabe M, Buth JE, Vishlaghi N, Torre-Ubieta L de la, Taxidis J, Khakh BS, et al. Self-Organized Cerebral Organoids with Human-Specific Features Predict Effective Drugs to Combat Zika Virus Infection. Cell Reports. 2017;21: 517–532. doi:10.1016/j.celrep.2017.09.047

31. Mulder LA, Vieira da Sá R, Korsten J, Freeze E, Schotting AJ, Koen G, et al. Effect of antivirals on clinical and lab-adapted human cytomegalovirus strains using induced pluripotent stem cell-derived human neural models. Antiviral Research. 2025;241: 106233. doi:10.1016/j.antiviral.2025.106233

32. Rozman B, Nachshon A, Levi Samia R, Lavi M, Schwartz M, Stern-Ginossar N. Temporal dynamics of HCMV gene expression in lytic and latent infections. Cell Rep. 2022;39: 110653. doi:10.1016/j.celrep.2022.110653

33. Gabrielli L, Bonasoni MP, Santini D, Piccirilli G, Chiereghin A, Petrisli E, et al. Congenital cytomegalovirus infection: patterns of fetal brain damage. Clinical Microbiology and Infection. 2012;18: E419–E427. doi:10.1111/j.1469-0691.2012.03983.x

34. Gale SD, Farrer TJ, Erbstoesser R, MacLean S, Hedges DW. Human Cytomegalovirus Infection and Neurocognitive and Neuropsychiatric Health. Pathogens. 2024;13: 417. doi:10.3390/pathogens13050417

35. Su X, Yue P, Kong J, Xu X, Zhang Y, Cao W, et al. Human Brain Organoids as an In Vitro Model System of Viral Infectious Diseases. Front Immunol. 2022;12. doi:10.3389/fimmu.2021.792316

36. Sun K, Fournier M, Sundberg AK, Song IH. Maribavir: Mechanism of action, clinical, and translational science. Clinical and Translational Science. 2024;17: e13696. doi:10.1111/cts.13696

37. Chou S, Winston DJ, Avery RK, Cordonnier C, Duarte RF, Haider S, et al. Comparative Emergence of Maribavir and Ganciclovir Resistance in a Randomized Phase 3 Clinical Trial for Treatment of Cytomegalovirus Infection. J Infect Dis. 2025;231: e470–e477. doi:10.1093/infdis/jiae469

38. Koszalka GW, Johnson NW, Good SS, Boyd L, Chamberlain SC, Townsend LB, et al. Preclinical and Toxicology Studies of 1263W94, a Potent and Selective Inhibitor of Human Cytomegalovirus Replication. Antimicrobial Agents and Chemotherapy. 2002;46: 2373–2380. doi:10.1128/aac.46.8.2373-2380.2002

39. Halpern-Cohen V, Blumberg EA. New Perspectives on Antimicrobial Agents: Maribavir. Antimicrobial Agents and Chemotherapy. 2022;66: e02405–21. doi:10.1128/aac.02405-21

40. Cheng MP, Gonzalez-Bocco IH, Arbonna-Haddad E, Aleissa M, Chen K, Zhou E, et al. Letermovir Treatment for Refractory or Resistant Cytomegalovirus Infection or Disease or with Concurrent Organ Dysfunction: A Phase 2 Open Label Study. J Assoc Med Microbiol Infect Dis Can. 2025;10: 6–14. doi:10.3138/jammi-2024-0016

41. Gabanti E, Borsani O, Colombo AA, Zavaglio F, Binaschi L, Caldera D, et al. Human Cytomegalovirus-Specific T-Cell Reconstitution and Late-Onset Cytomegalovirus Infection in Hematopoietic Stem Cell Transplantation Recipients following Letermovir Prophylaxis. Transplantation and Cellular Therapy. 2022;28: 211.e1–211.e9. doi:10.1016/j.jtct.2022.01.008

42. Combs JA, Norton EB, Saifudeen ZR, Bentrup KHZ, Katakam PV, Morris CA, et al. Human Cytomegalovirus Alters Host Cell Mitochondrial Function during Acute Infection. Journal of Virology. 2020;94: 10.1128/jvi.01183-19. doi:10.1128/jvi.01183-19

43. Karniely S, Weekes MP, Antrobus R, Rorbach J, van Haute L, Umrania Y, et al. Human Cytomegalovirus Infection Upregulates the Mitochondrial Transcription and Translation Machineries. mBio. 2016;7: 10.1128/mbio.00029-16. doi:10.1128/mbio.00029-16

44. Yu Y, Clippinger AJ, Alwine JC. Viral effects on metabolism: changes in glucose and glutamine utilization during human cytomegalovirus infection. Trends in Microbiology. 2011;19: 360–367. doi:10.1016/j.tim.2011.04.002

45. Chen H, Beardsley GP, Coen DM. Mechanism of ganciclovir-induced chain termination revealed by resistant viral polymerase mutants with reduced exonuclease activity. Proceedings of the National Academy of Sciences. 2014;111: 17462–17467. doi:10.1073/pnas.1405981111

46. Prichard MN. Function of human cytomegalovirus UL97 kinase in viral infection and its inhibition by maribavir. Reviews in Medical Virology. 2009;19: 215–229. doi:10.1002/rmv.615

47. Acosta E, Bowlin T, Brooks J, Chiang L, Hussein I, Kimberlin D, et al. Advances in the Development of Therapeutics for Cytomegalovirus Infections. J Infect Dis. 2020;221: S32– S44. doi:10.1093/infdis/jiz493

48. Mei M-J, Zhou Y-P, Pan Y-T, Sun J-Y, Zeng W-B, Wu T, et al. Molecular features of congenital cytomegalovirus infection in neonatal mouse brain at single-cell resolution. acta neuropathol commun. 2025;13: 237. doi:10.1186/s40478-025-02158-x

49. Lancaster MA, Knoblich JA. Generation of cerebral organoids from human pluripotent stem cells. Nat Protoc. 2014;9: 2329–2340. doi:10.1038/nprot.2014.158

50. Hamilton ST, Marschall M, Rawlinson WD. Investigational Antiviral Therapy Models for the Prevention and Treatment of Congenital Cytomegalovirus Infection during Pregnancy. Antimicrob Agents Chemother. 2020;65: e01627–20. doi:10.1128/AAC.01627-20

51. Hamilton ST, Hutterer C, Egilmezer E, Steingruber M, Milbradt J, Marschall M, et al. Human cytomegalovirus utilises cellular dual-specificity tyrosine phosphorylation-regulated kinases during placental replication. Placenta. 2018;72–73: 10–19. doi:10.1016/j.placenta.2018.10.002

52. Hutterer C, Hamilton S, Steingruber M, Zeitträger I, Bahsi H, Thuma N, et al. The chemical class of quinazoline compounds provides a core structure for the design of anticytomegaloviral kinase inhibitors. Antiviral Research. 2016;134: 130–143. doi:10.1016/j.antiviral.2016.08.005

53. Patel H, Manning J, Ewels P, Garcia MU, Peltzer A, Hammarén R, et al. nf-core/rnaseq: nf-core/rnaseq v3.21.0 - Mercury Macaw. Zenodo; 2025. doi:10.5281/zenodo.17153746

54. da Veiga Leprevost F, Grüning BA, Alves Aflitos S, Röst HL, Uszkoreit J, Barsnes H, et al. BioContainers: an open-source and community-driven framework for software standardization. Bioinformatics. 2017;33: 2580–2582. doi:10.1093/bioinformatics/btx192

55. Andrews S. FastQC: a quality control tool for high throughput sequence data. 2010;Available from: http://www.bioinformatics.babraham.ac.uk/projects/fastqc.

56. Chen S, Zhou Y, Chen Y, Gu J. fastp: an ultra-fast all-in-one FASTQ preprocessor. Bioinformatics. 2018;34: i884–i890. doi:10.1093/bioinformatics/bty560

57. Patro R, Duggal G, Love MI, Irizarry RA, Kingsford C. Salmon provides fast and bias-aware quantification of transcript expression. Nature methods. 2017;14: 417.

58. Kanton S, Boyle MJ, He Z, Santel M, Weigert A, Sanchís-Calleja F, et al. Organoid single-cell genomic atlas uncovers human-specific features of brain development. Nature. 2019;574: 418–422. doi:10.1038/s41586-019-1654-9

59. Jew B, Alvarez M, Rahmani E, Miao Z, Ko A, Garske KM, et al. Accurate estimation of cell composition in bulk expression through robust integration of single-cell information. Nat Commun. 2020;11: 1971. doi:10.1038/s41467-020-15816-6

60. Love MI, Soneson C, Hickey PF, Johnson LK, Pierce NT, Shepherd L, et al. Tximeta: Reference sequence checksums for provenance identification in RNA-seq. PLOS Computational Biology. 2020;16: e1007664. doi:10.1371/journal.pcbi.1007664

61. Chen Y, Chen L, Lun ATL, Baldoni PL, Smyth GK. edgeR v4: powerful differential analysis of sequencing data with expanded functionality and improved support for small counts and larger datasets. Nucleic Acids Res. 2025;53: gkaf018. doi:10.1093/nar/gkaf018

62. Ritchie ME, Phipson B, Wu D, Hu Y, Law CW, Shi W, et al. limma powers differential expression analyses for RNA-sequencing and microarray studies. Nucleic Acids Res. 2015;43: e47. doi:10.1093/nar/gkv007

63. Wu D, Smyth GK. Camera: a competitive gene set test accounting for inter-gene correlation. Nucleic Acids Res. 2012;40: e133. doi:10.1093/nar/gks461

64. Liberzon A, Birger C, Thorvaldsdóttir H, Ghandi M, Mesirov JP, Tamayo P. The Molecular Signatures Database Hallmark Gene Set Collection. cels. 2015;1: 417–425. doi:10.1016/j.cels.2015.12.004

65. Gu Z. Complex heatmap visualization. iMeta. 2022;1: e43. doi:10.1002/imt2.43

66. Wickham H. ggplot2. WIREs Computational Statistics. 2011;3: 180–185. doi:10.1002/wics.147

67. Hall MB, Coin LJM. Pangenome databases improve host removal and mycobacteria classification from clinical metagenomic data. Gigascience. 2024;13: giae010. doi:10.1093/gigascience/giae010

68. Shaw J, Yu YW. Rapid species-level metagenome profiling and containment estimation with sylph. Nat Biotechnol. 2025;43: 1348–1359. doi:10.1038/s41587-024-02412-y

69. Foster CSP, Walker GJ, Jean T, Wong M, Brassil L, Isaacs SR, et al. Long-term serial passaging of SARS-CoV-2 reveals signatures of convergent evolution. Journal of Virology. 2025;99: e00363–25. doi:10.1128/jvi.00363-25

70. Foster CSP, Rawlinson WD. vartracker: an end-to-end tool for pathogen longitudinal variant analysis and visualisation. bioRxiv; 2026. p. 2026.05.06.723370. doi:10.64898/2026.05.06.723370

